# HilE represses the activity of HilD via a mechanism distinct from that of intestinal long-chain fatty acids

**DOI:** 10.1101/2023.02.23.529715

**Authors:** Joe D. Joiner, Wieland Steinchen, Thales Kronenberger, Gert Bange, Antti Poso, Samuel Wagner, Marcus D. Hartmann

## Abstract

The expression of virulence factors essential for the invasion of host cells by *Salmonella enterica* is tightly controlled by a network of transcription regulators. The AraC/XylS transcription factor HilD is the main integration point of environmental signals into this regulatory network, with many factors affecting HilD activity. Long chain fatty acids (LCFAs), which are highly abundant throughout the host intestine directly bind to, and repress HilD, acting as environmental cues to coordinate virulence gene expression. The regulatory protein HilE also negatively regulates HilD activity, through a protein-protein interaction. Both of these regulators inhibit HilD dimerisation, preventing HilD from binding to target DNA. We investigated the structural basis of these mechanisms of HilD repression. LCFAs bind to a conserved pocket in HilD, in a comparable manner to that reported for other AraC/XylS regulators, whereas HilE forms a stable heterodimer with HilD by binding to the HilD dimerisation interface. Our results highlight two distinct mechanisms by which HilD activity is repressed, which could be exploited for the development of new antivirulence leads.

## Introduction

*Salmonella enterica* is an enteric pathogen and one of the leading causes of gastrointestinal disease. *Salmonella spp.* adhere to and invade epithelial cells via a complex mechanism requiring many virulence factors, most of which are located on five highly-conserved horizontally-acquired *Salmonella* pathogenicity islands (SPIs)[1–3]. *Salmonella* pathogenicity island 1 (SPI-1) encodes the genes required for the initial invasion of host cells, including numerous effector proteins and a type III secretion system (T3SS-1) injectosome that enables the direct injection of proteins from the bacterial cytoplasm into the host cell[4,5]. These effectors have a range of purposes, including causing changes in the host cell actin cytoskeleton that results in the engulfment of *Salmonella* cells by endocytosis.

To coordinate the sequential expression of different virulence genes according to the stage of the infection process, expression of SPI genes is tightly regulated. The transcriptional regulator HilA activates the expression of the *prg/org* and *inv/spa* operons, which encode the structural components of the T3SS-1, and the genes encoding several of the effector proteins secreted through T3SS-1[6–8]. Expression of *hilA* is in turn controlled by the action of three AraC/XylS transcription regulators: HilD, HilC and RtsA, which bind to overlapping sites within the *hilA* promoter to active expression[9,10]. The AraC/XylS family is defined by a highly conserved DNA-binding domain (DBD) containing two helix-turn-helix (HTH) motifs, which form direct contacts with DNA[11–13]. HilD, HilC and RtsA all have a two-domain structure comprising an N-terminal regulatory domain and a C-terminal DBD, which is the most common domain organisation of AraC/XylS family proteins[13]; examples include the regulators ToxT and Rns, for which full-length structures have been experimentally determined[14,15]. Each of HilD/HilC/RtsA is able to activate not only *hilA*, but also its own promoter and that of the other two regulators, forming a complex feed-forward loop to activate SPI-1 expression (Fig 1)[16]. HilD is the most prominent activator of *hilA*, with HilC and RtsA serving to amplify *hilA* transcription[16,17].

**Fig 1.**
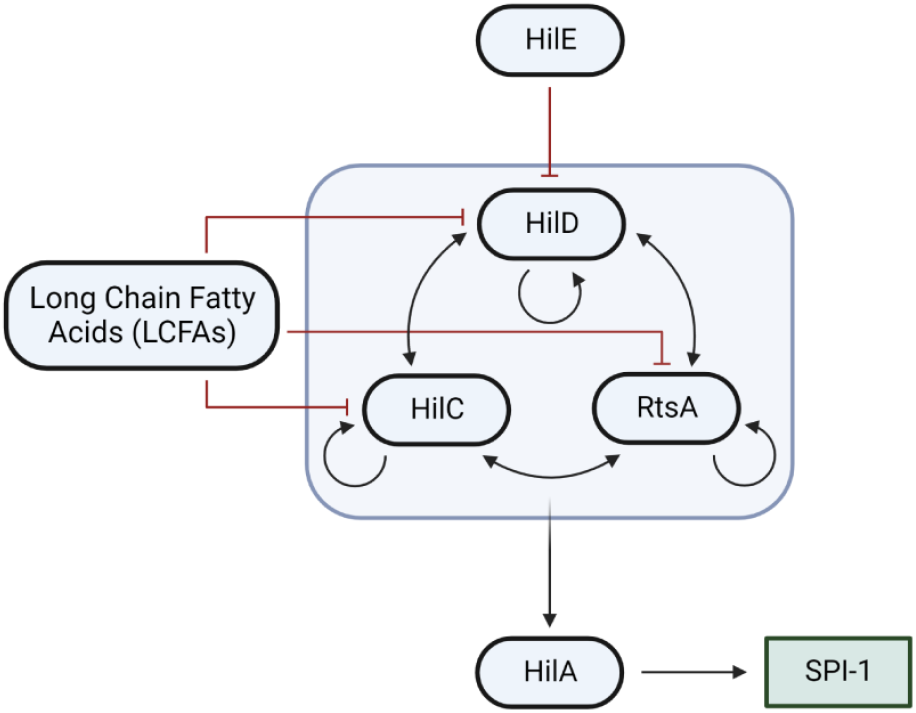
Simplified model of the SPI-1 regulatory network. Black arrows indicate activation and red lines with blunt ends represent repression.

HilD is additionally the main integration point of environmental signals into the SPI-1 regulatory network, as many regulatory factors affect *hilD* transcription, translation, or HilD activity to regulate *hilA* activation[18]. HilD/HilC/RtsA can bind a range of small molecules, which regulate their ability to bind to target DNA. These include long chain fatty acids (LCFAs) and bile acids present in the gut, which *Salmonella* utilise to sense their intestinal location and coordinate the expression of virulence genes at specific locations where invasion can occur[19–22]. Other compounds have also been shown to bind to these regulators to inhibit *hilA* expression and *Salmonella* invasion, highlighting the potential of these regulators as targets of novel antipathogenic compounds[23,24].

One of the most important negative regulators of HilD is the regulatory protein HilE. HilE is a homolog of hemolysin-coregulated protein (Hcp), a key structural component of the Type VI Secretion System (T6SS), and specifically represses HilD activity through a protein-protein interaction[25–27]. However, the mechanism and structural basis of this interaction remain elusive.

Here we used a range of biochemical and biophysical methods to elucidate and compare the regulation of the HilD protein by LCFAs and HilE. We show that LCFAs bind to, and regulate, HilD in a comparable mechanism to other AraC/XylS regulators of virulence genes. HilE forms a stable heterodimer with HilD, disrupting HilD homodimerisation and preventing HilD from binding to target DNA. Our results highlight the different biochemical mechanisms that to repress HilD activity, and a unique mechanism for the regulation of AraC/XylS transcription factors.

## Results

### Structural and dimerisation characteristics of HilD

The structural characterisation of AraC/XylS proteins is challenging due to their poor solubility at higher concentrations[28], and an experimental structure of HilD remains elusive. The predicted structure of HilD, retrieved from the AlphaFold database (AlphaFold EBI ID P0CL08), concurs with the expected domain organisation (Fig 2A) and is comparable to other AraC/XylS proteins for which full-length structures have been experimentally determined[14,15,29]. The N-terminal domain (NTD) contains a cupin barrel structure, which has been shown to form the fatty acid binding site in the AraC/XylS family members ToxT and Rns, as well as a number of helices that form the reported dimerisation interface. The C-terminal DNA-binding domain is comprised of 7 α-helices (α7-α13), which constitute two HTH motifs connected by an α-helical linker (helix α10). The N-terminus of HilD (residues 1-35) was predicted with very low confidence (pLDDT < 50%) in the AlphaFold model indicative of disorder, and hence is hidden in all protein figures for clarity.

**Fig 2.**
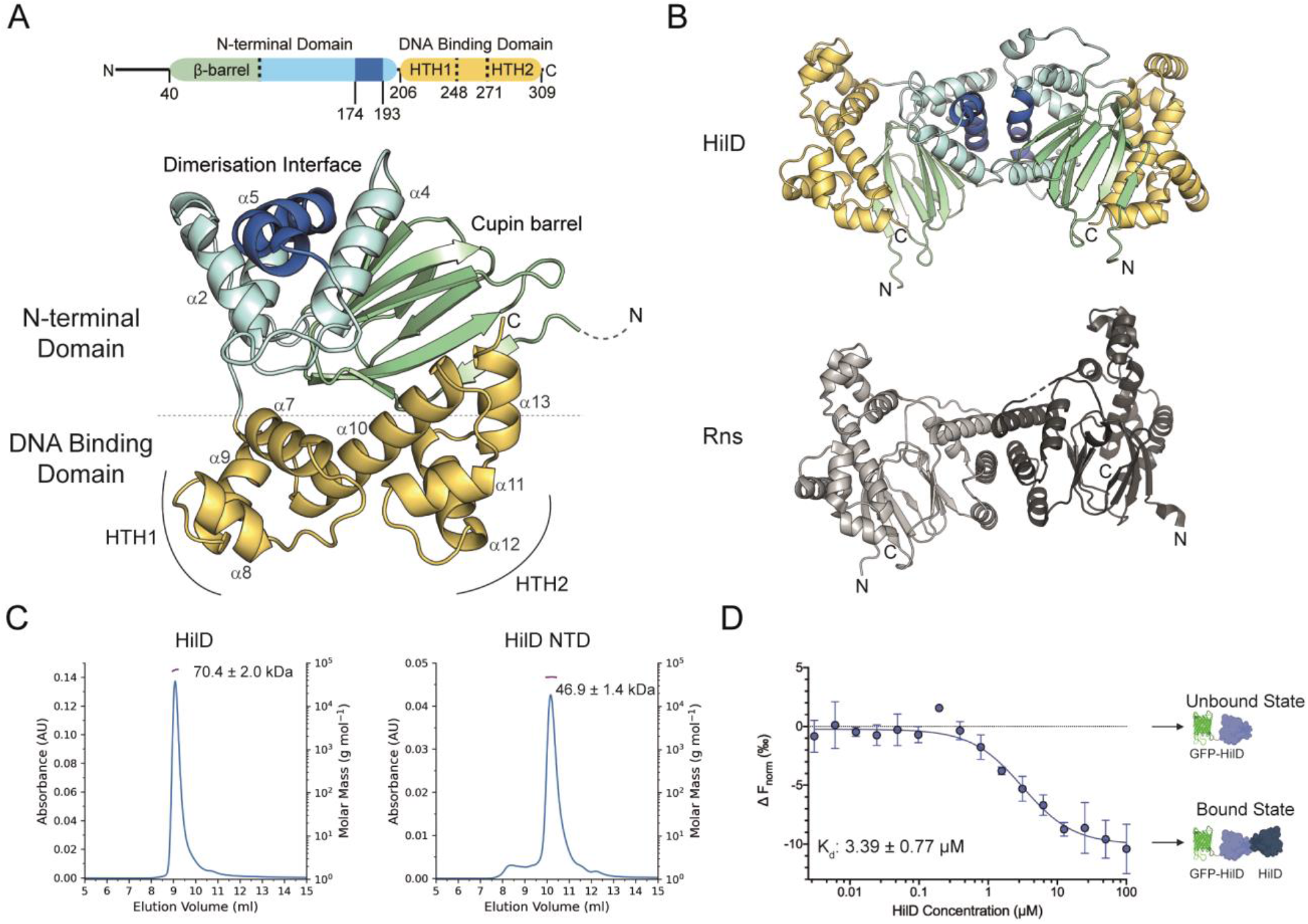
HilD forms homodimers reminiscent of other AraC/XylS transcription regulators. (**A**) AlphaFold2 model of HilD. The N-terminal domain is coloured in cyan, with the cupin barrel and reported dimerisation helix highlighted in green and dark blue, respectively. The DNA binding domain is coloured in yellow, with the helices constituting this domain and the two HTH motifs labelled. (**B**) AlphaFold2 model of the HilD homodimer (top) and the crystal structure of the Rns homodimer (PDB: 6XIV) (bottom). HilD residues 1-35 are removed in both (A) and (B) clarity. (**C**) SEC-MALS profiles of full-length and the N-terminal domain (NTD) of HilD. Calculated molecular weight values correspond to 3 repeat experiments. (**D**) HilD dimerisation measured by MST. Unlabelled HilD protein (3.05 nM to 100 μM) was incubated with 50 nM GFP-HilD.

HilD is known to form both homodimers and heterodimers with the other SPI-1 regulators HilC and RtsA[30], and we modelled the HilD homodimer using AlphaFold multimer[31]. The predicted homodimer topology of HilD is comparable to that of the AraC/XylS proteins Rns and ExsA for which the dimeric structure has been experimentally determined[15,32] (Fig 2B). We used multi-angle light scattering coupled to size-exclusion chromatography (SEC-MALS) to confirm that our purified full length HilD exists exclusively as a dimer in solution (Fig 2C). We also purified a construct lacking the DNA binding domain (HilD NTD, residues 7-206) that similar to full-length HilD appears dimeric during SEC-MALS runs (Fig 2C), in agreement with previous data showing that the NTD is responsible for dimerisation, which is primarily mediated by helix α5, formed by residues 180-192, at the centre of the dimerisation interface[30].

We conceived a microscale thermophoresis (MST) assay to quantify the homodimerization of HilD, in which HilD was fused to an N-terminal GFP tag for detection. GFP-HilD, at a constant concentration of 50 nM, was then incubated with varying concentrations of unfused HilD. A dose-dependent reduction in normalized fluorescence yielded an equilibrium dissociation constant for HilD dimerisation, K_d,dimer_, of 3.39 ± 0.77 μM (Fig 2D). Collectively, these data suggest that HilD exhibits the typical characteristics of AraC/XylS protein family members.

### HilD Binds a Range of Fatty Acids

HilD is capable of binding a range of different LCFAs, and a number of residues were previously reported to contribute to this interaction[20]. Using MST, we first determined the affinity of oleic acid, which has previously been shown to bind to HilD to repress *hilA* expression[19,20,33], to HilD and obtained a K_d_ of 48 ± 5.27 μM for this interaction (Fig 3A).

**Fig 3.**
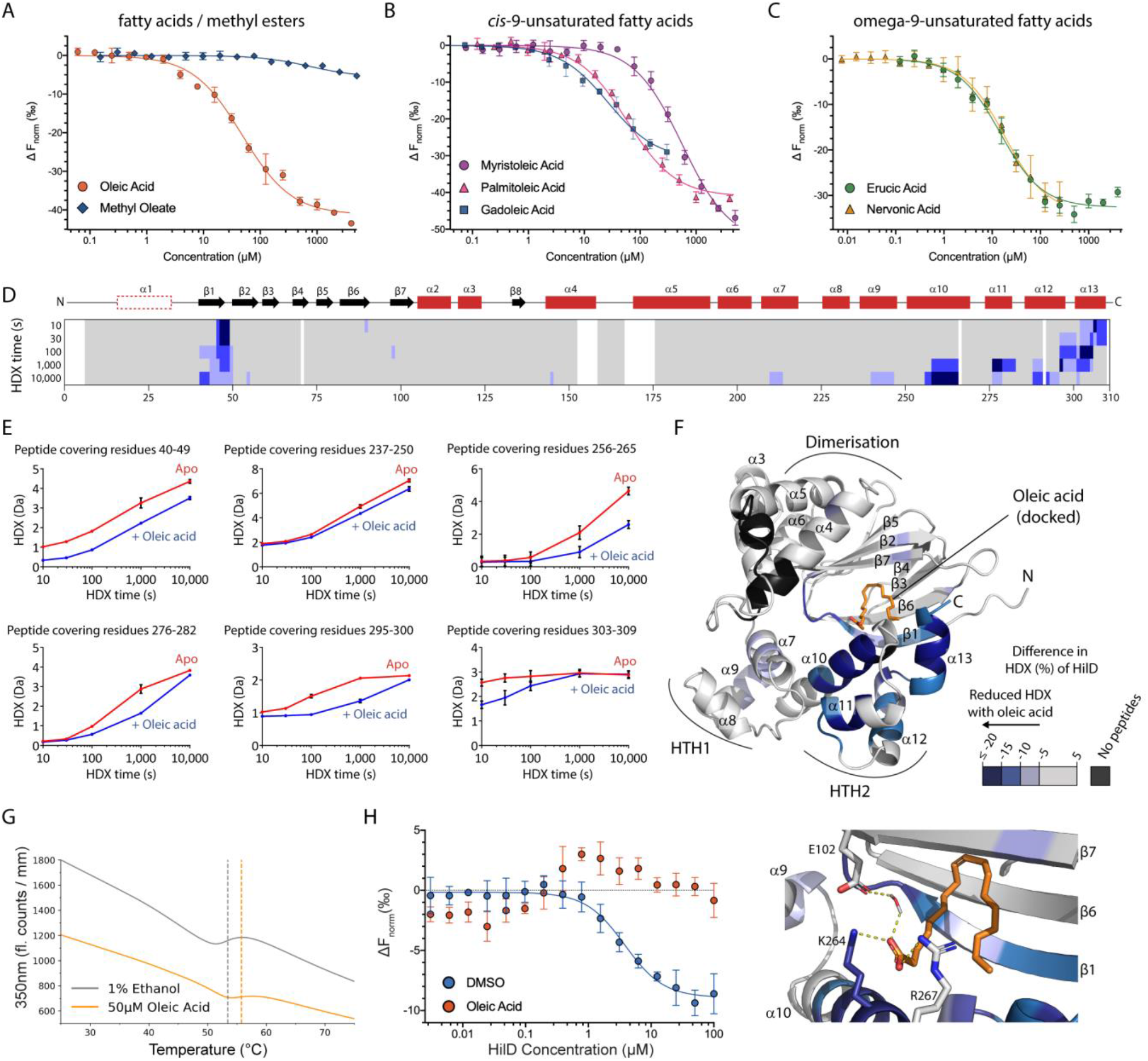
Long chain fatty acids bind to a conserved binding pocket in HilD. (**A-C**) MST binding curves for fatty acid binding to HilD: (A) oleic acid and methyl oleate; (B) *cis*-9-unsaturated fatty acids; (C) omega-9-unsaturated fatty acids. Calculated binding affinities are displayed in Table 1. (**D**) The difference in HDX between oleic acid-bound and apo HilD projected on its amino acid sequence. Different tones of blue reflect reduction in HDX of HilD in presence of oleic acid. The HilD secondary structure is schematically depicted above. (**E**) Representative HilD peptides displaying changes in HDX with respect to oleic acid. Data represent the mean ± s.d. of n=3 replicates. (**F**) The oleic acid-dependent HDX changes were projected onto the structural model of HilD with the oleic acid binding site inferred by molecular docking of the ligand. In the detailed view of the oleic acid binding pocket, residues predicted to form specific interactions with oleic acid are displayed as sticks, with hydrogen bonds shown as yellow dashes. (**G**) NanoDSF unfolding profiles of HilD (20 μM) when incubated with either 50 μM oleic acid or 1% ethanol. Traces show the average fluorescent readout obtained from 10 technical replicates. The HilD melting temperature calculated for both samples is indicated by the dashed vertical lines. (**H**) MST dose-response plots for the homodimerisation of HilD, whereby GFP-HilD was pre-incubated with 1% DMSO (blue), or 100 μM oleic acid (orange). MST experiments were carried out in triplicate and the K_d,dimer_ was determined from changes in thermophoresis at an MST on-time of 1.5 seconds.

We then determined the binding affinity of a number of other unsaturated fatty acids, to establish which structure-related properties of these ligands are critical for binding to HilD. We first examined the effect of varying the chain length of LCFAs containing a *cis-9* double bond, as in oleic acid. Binding affinity increased with increasing chain length (Fig 3B). Myristoleic acid (C14) displays only very weak binding (Kd of 567 μM), signifying 16 carbon atoms as the minimum chain length of FAs that is required for efficient binding to HilD. The same trend of higher affinity for longer chain unsaturated fatty acids was observed with increasing chain length between the *cis*- double bond and the carboxylic acid head group (Fig 3C). These combined results imply that the position of this central double bond is not a requirement for binding, supported by previous findings that *cis*-2-unsaturated fatty acids, which lack a double bond in the centre of the hydrocarbon chain, bind to HilD with high affinity[33]. The comparable affinity of erucic acid (C22, 14.71 ± 2.19 μM) and nervonic acid (C24, 17.99 ± 1.84 μM) suggests a maximum length of 22 carbon atoms is optimal for binding to HilD, and LCFAs of longer length may be protruding from the binding pocket. We also attempted to determine the affinity of corresponding *trans*-unsaturated and saturated LCFAs, however, the low solubility of these compounds prevented us from performing affinity measurements. To investigate whether specific interactions involving the carboxylic acid head group are important for LCFA binding, we compared the binding of oleic acid with the corresponding methyl ester (Fig 3A). These methyl esters did not bind to HilD, in agreement with a previous study showing methyl esters of *cis*-2-UFAs showed reduced potency in repressing *hilA*[33].

### LCFAs bind to HilD in a comparable manner to other AraC/XylS proteins

To experimentally probe the LCFA binding pocket in HilD, we used hydrogen-deuterium exchange mass spectrometry (HDX-MS). HDX-MS detects changes in the accessibility of backbone amide hydrogens, which undergo exchange with deuterium in deuterated buffers on a time scale measurable by MS, and thus provides snapshots of a protein’s higher order structure and changes therein by e.g., ligand binding. HDX-MS was performed on HilD in both the absence and presence of oleic acid, which was previously shown to bind to HilD to repress *hilA* expression[19,20,33]. We mapped detected changes in HDX upon incubation with oleic acid onto the predicted structure of HilD to identity the location of the binding pocket (Fig 3D-F). Decreased HDX was observed over the entire DNA binding domain, but most pronounced at the C-terminal portion of helix α10 and helices α11-13, and β-strand 1, which altogether line the cavity provided by the β-barrel (Fig 3F). Only minor changes were apparent for the other β-strands forming the barrel structure, which may be reasoned by the intrinsically very low HDX rate of these entities (Dataset S4). Mildly altered HDX in α7 and α9 (HTH1) may be a consequence of the change in α10, preventing the independent rotation of the two HTH motifs with respect to one another. Collectively, the vicinity of the strongest HDX decreases suggests the β-barrel/HTH2 interface as the oleic acid binding pocket in agreement with the fatty acid binding pocket in ToxT and Rns[14,15], and computational docking of oleic acid to HilD (Fig 3F). In this model, specific polar interactions are predicted between the carboxyl head group of oleic acid with residues K264 and R267, which are located on the DBD within a region of decreased HDX, and water-mediated interactions with residue E102. (Fig 3F). We also found that the binding of oleic acid increased the melting temperature of HilD (Fig 3G), similar to the effect observed for fatty acid binding to Rns[15]. This is consistent with our model showing that oleic acid forms specific interactions with residues on both domains, confining HilD to a more rigid structure.

The binding of small molecules has previously been shown to disrupt the dimerisation of both HilD[22] and ToxT[34], and we used the MST dimerisation assay (Fig 2D) to investigate the effects of oleic acid on HilD homodimerisation. Whilst DMSO had no significant effect on HilD dimerisation (K_d,dimer_: 4.52 ± 0.50 μM), when GFP-HilD was incubated with oleic acid (100 μM), the formation of heterodimers between GFP-HilD and unfused HilD was completely abolished (Fig 3H). This shows that LCFAs disrupt the dimerisation of HilD to prevent HilD from binding to target DNA, in a similar manner as reported for ToxT[35,36].

### HilE Does Not Form Higher Order Oligomers

An experimentally determined structure of HilE remains elusive, however, its predicted structure is comparable to that of Hcp family proteins, comprising a tight β-barrel domain with a single α-helix (residues 58-68) located on one side of the β-barrel (Fig S1D). Although the C-terminus is predicted with low confidence in the AlphaFold model, indicative of disorder, other prediction servers modelled the C-terminus as an additional β-strand, as observed in the structures of other Hcp proteins (Fig S1D).

Other Hcp family proteins assemble into hexameric ring structures. Based on its predicted structural similarity to these proteins, HilE is also postulated to form oligomers and it was previously reported that HilE may inhibit HilD activity through the formation of a large protein complex with HilD[26]. We recombinantly expressed and purified HilE to homogeneity in high yields (Fig S1A) and determined its oligomerisation state using SEC-MALS. Surprisingly, HilE was found to exist exclusively as monomers in solution (Fig S1C). We expressed several different HilE constructs with different purification tags confirming that the additional residues introduced by tags at either terminus were not inhibiting ring formation (Fig S2). An extended loop region, which can be defined from the structural alignment of Hcp proteins, has previously been shown to be crucial for the assembly of Hcp proteins into hexameric rings and subsequent ring stacking during nanotube formation[37]. Mutants of *Salmonella* Hcp2, containing mutations within this loop region were defective in ring formation[38]. This loop is also notably shorter in HilE (formed by residues T34-Y45) than in other Hcp-like proteins (Figs S1D and S1E), explaining the observation that HilE does not appear to form hexameric rings.

### HilE Forms a Stable Heterodimer with HilD

HilE has previously been shown to prevent HilD from binding to target DNA, however, the mechanism of this inhibition is elusive and whether HilE also affects HilD dimerisation has been subject to debate[26,27]. We found that HilD dimerisation was no longer observed in our MST-based dimerisation assay when GFP-HilD was first incubated with excess HilE (Fig 4A), akin to the effect seen for oleic acid. This suggests that HilE negatively regulates HilD activity by inhibiting HilD dimerisation and subsequently preventing DNA binding. We determined the affinity of the HilD-HilE interaction by MST, showing that HilE binds to HilD with a K_d_ of 1.68 ± 0.47 μM (Fig 4B). To further characterise the interaction between HilD and HilE, we next performed a gel filtration assay to confirm the two proteins were able to form a stable complex. A 1:1 mixture of the two proteins resulted in a single elution peak, corresponding to the heterocomplex (Fig 4C).

**Fig 4.**
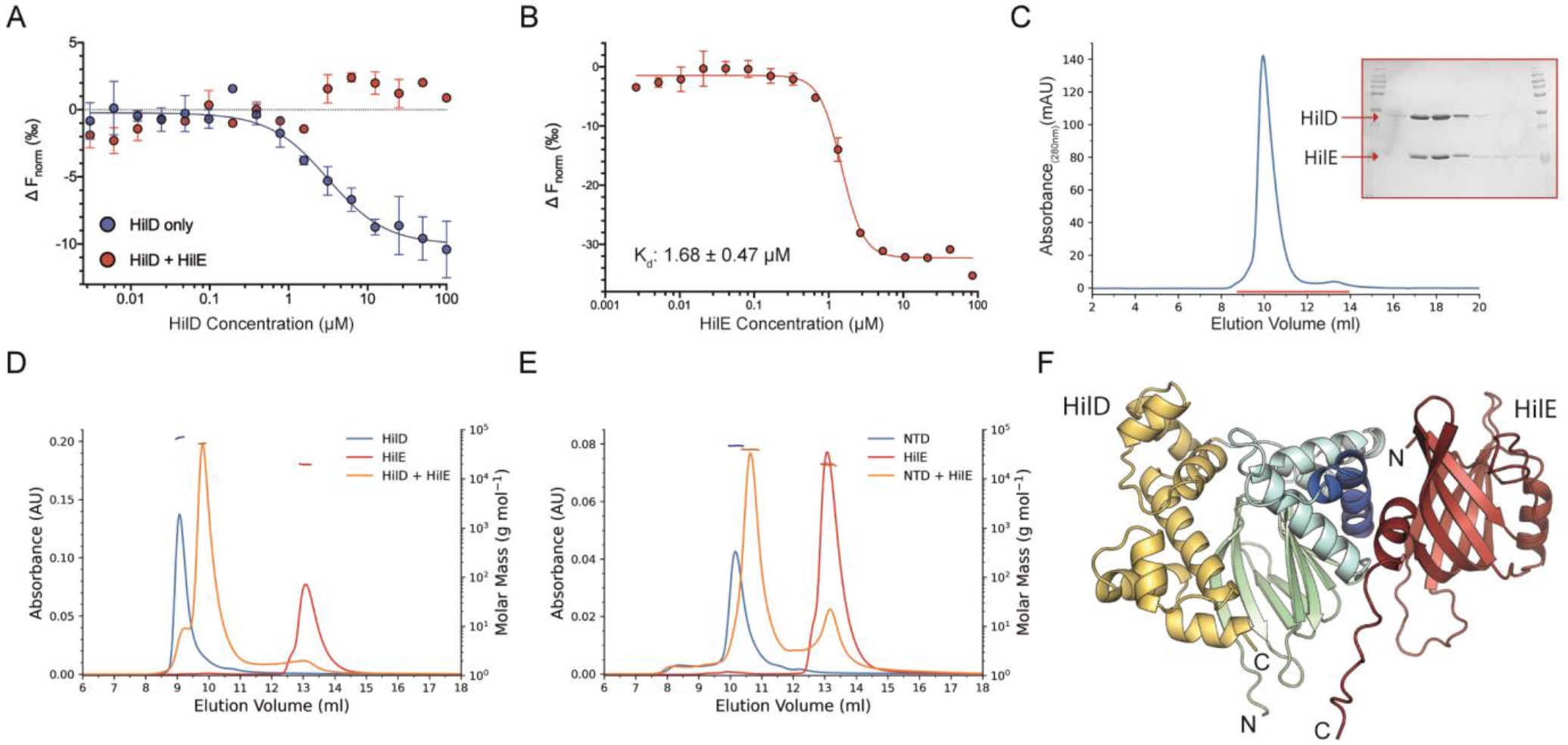
HilD and HilE form a stable heterodimer. (**A**) Homodimerisation of HilD monitored by MST. Unlabelled HilD protein (3.05 nM to 100 μM) was incubated with 50 nM GFP-HilD, in the presence (red) or absence (blue) of 10 μM HilE. K_d,dimer_ was determined from changes in thermophoresis at an MST on-time of 1.5 seconds, with error bars showing standard deviation from three repeat experiments. (**B**) MST dose-response curve for the binding of HilD and HilE. HilE was titrated against GFP-HilD (50 nM) and the K_d_ calculated from changes in thermophoresis at 2.5 seconds on-time (three replicates). (**C**) Elution profile for the purification of the HilD/HilE complex using a S75 10/300 increase size exclusion chromatography column. Selected fractions, highlighted by the red line in the elution trace, were loaded to an SDS gel and stained with Coomassie. (**D-E**) SEC-MALS analysis of (D) HilD and HilE and (E) HilD NTD and HilE. Left x-axis shows UV absorbance measured at 280nm; right x-axis shows the calculated molecular weight values from light scattering, highlighted by horizontal dashes, with values displayed in Table 2. (**F**) Highest ranked model of the HilD-HilE heterodimer, predicted using AlphaFold Multimer. HilD is coloured as in Fig 1, with HilE coloured in red.

We determined the molecular mass of the HilD/HilE complex to be 52.7 ± 0.9 kDa using SEC-MALS, confirming the formation of a heterodimer (Fig 4D). This is consistent with the observation that HilE disrupts HilD dimerisation, as both shown by our MST assay and reported previously[27]. We performed additional SEC-MALS runs using the HilD NTD construct, rather than full-length HilD. The formation of a stable heterodimer was again observed, showing that HilE interacts with the HilD NTD and that the HilD DBD is not required for binding (Fig 4E). We also purified the other *hilA* activator HilC (Fig S3A-C) and performed SEC-MALS analysis, to investigate any potential interaction between HilC and HilE. No complex formation between HilC and HilE was observed, with two clear peaks in the UV trace corresponding to the elution of the individual proteins (Fig S3E). These results show that HilE interacts specifically with the NTD of HilD, in line with a previous study showing that HilE inhibits the DNA-binding activity of HilD, but not that of HilC[26].

### HilE may directly replace one of the monomers of the HilD dimer

We modelled the HilD-HilE heterodimer complex using AlphaFold multimer[30], with the predicted structure showing that HilE directly disrupts HilD dimerisation by displacing one of the two HilD molecules constituting the dimer pair (Fig 4F). In all predicted models, the dimerisation helix of HilD binds to the opposite face of the HilE β-barrel to that of the HilE α-helix, similar to the interactions between neighbouring subunits in the hexameric structures formed by other Hcp proteins. In both the HilD homodimer and HilD-HilE heterodimer, the interface is dominated by hydrophobic interactions, however, the predicted HilD-HilE binding interface also contains several hydrogen bond and salt bridge interactions. We determined the residues forming the HilD homodimer interface using PISA[39], and calculated the buried surface area to be > 935 Å^2^ in all predicted models. For the HilD-HilE complex prediction, the calculated buried surface area lies between 518-779 Å^2^ for the 10 highest ranked models, suggesting that the binding interface of the HilD-HilE heterodimer may be smaller than that in the HilD homodimer.

To experimentally probe our model of HilD-HilD homodimer disruption by formation of the HilD-HilE heterodimer, we performed HDX-MS experiments for both individual HilD and HilE proteins and the HilD-HilE complex (Figs 5 and 6). Increased HDX covering most of the dimerisation interface of HilD in the context of the HilD-HilE heterodimer is consistent with the notion that HilE disrupts HilD homodimerisation (Fig 5A-C). No change though was apparent for residues 177-183 at the centre of the HilD dimerisation helix α5 (Fig 5) and a number of peptides covering helix α4 displaying perturbed HDX, characterized by an HDX increase after short HDX incubation times but decrease at longer HDX times (representative peptide covering residues 135-144, Fig 5B). Both α4 and α5 reflect the primary contact sites for HilE in our model of the HilD-HilE complex (Fig 5D) whereby the mixed HDX behaviour (both increased and decreased HDX) at the HilD dimerization and presumed HilD-HilE interface seems to reflect different binding modes and the differences in buried surface areas of the homodimeric and heterodimeric complexes.

**Fig 5.**
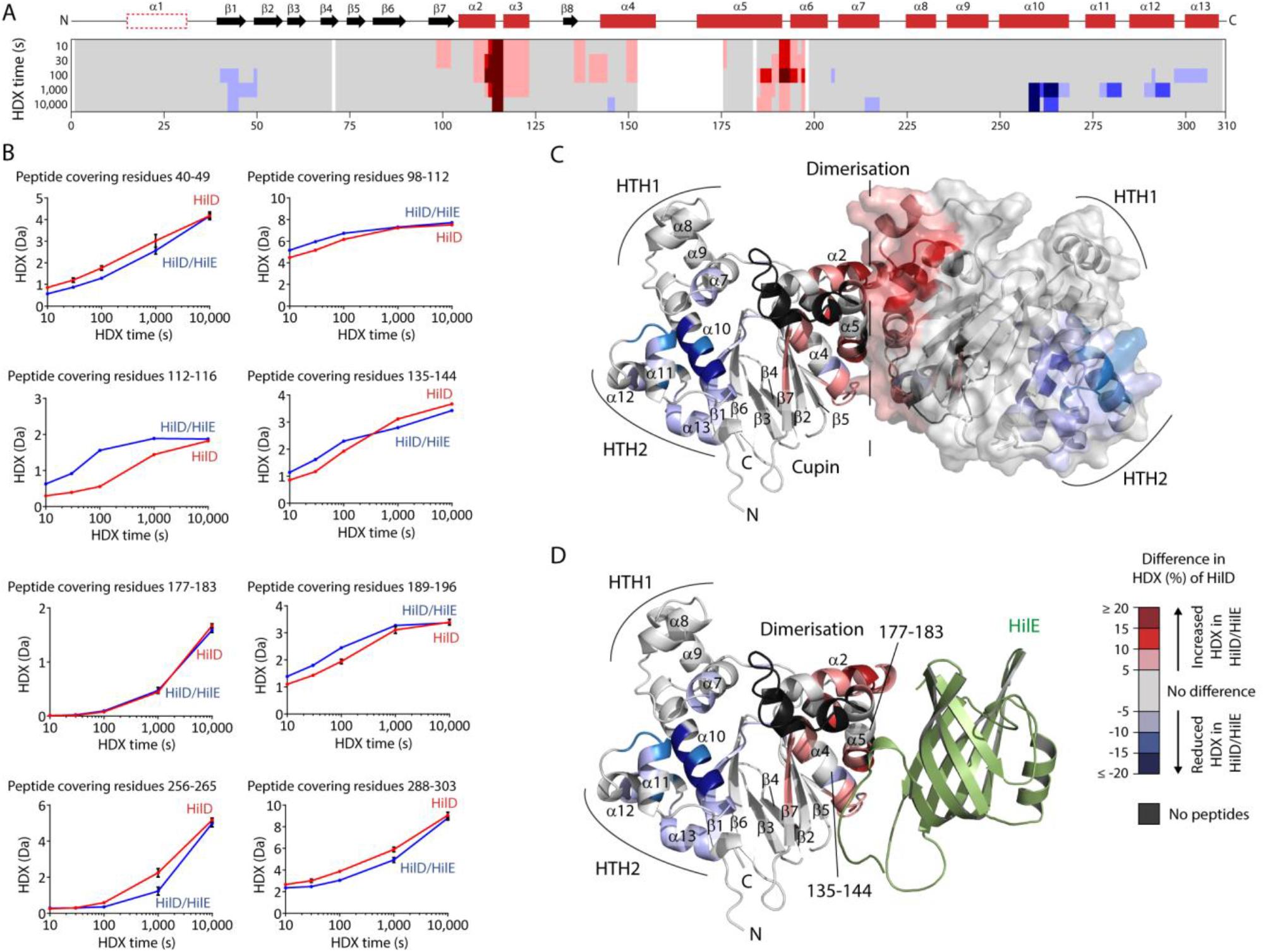
Conformational changes of HilD in the HilD/HilE complex. (**A**) The difference in HDX between the HilD-HilE complex and individual HilD projected onto the HilD amino acid sequence. Different tones of red and blue reflect increased and decreased HDX of HilD the complex. The HilD secondary structure is schematically depicted above. (**B**) Representative HilD peptides displaying changes in HDX. Data represent the mean ± s.d. of n=3 replicates. (**C-D**) The altered HDX of HilD in the HilD-HilE complex was projected onto a model of (**C**) the HilD-HilD homodimer, or (**D**) the HilD-HilE heterodimer.

**Fig 6.**
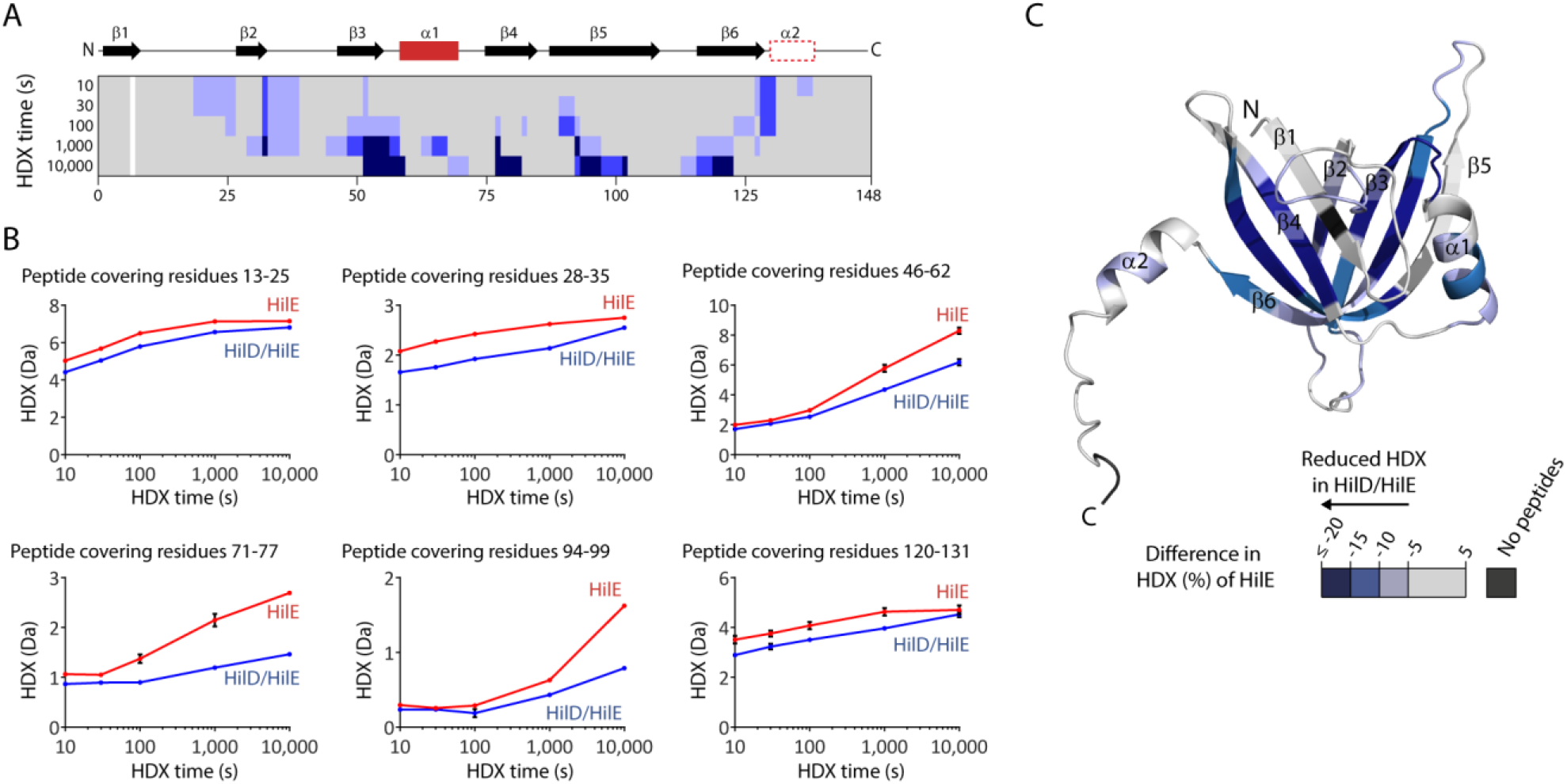
Conformational changes of HilE in the HilD/HilE complex. (**A**) The difference in HDX between the HilD-HilE complex and individual HilE projected onto the HilE amino acid sequence. Different tones of blue reflect decreased HDX of HilE when in complex with HilD. The HilE secondary structure is schematically depicted above. (**B**) Representative HilE peptides displaying changes in HDX. Data represent the mean ± s.d. of n=3 replicates. (**C**) The difference in HDX from (A) projected onto a model of HilE.

Decreased HDX of HilD upon HilE binding is observed at residues 41-50, the C-terminal end of helix α10 (residues 258-268) and helices α11-α13 of the DBD, which comprise HTH2 and the surrounding regions. Given that our SEC-MALS experiments suggested that the DBD is not required for the HilD/HilE interaction (Fig 4E), we hypothesize that these areas do not reflect the HilE binding site itself (or a part of it) on HilD but are more likely conformational changes associated with HilE binding. Notably, these areas with reduced HDX in HilD-HilE are reminiscent of the changes induced by oleic acid (Fig 3D-F), which like HilE impairs HilD DNA binding ability.

We also analysed the changes in HDX of HilE upon binding to HilD. Hereby, decreased HDX was apparent across the entire HilE domain and portions of the flanking helices (Fig 6), implying that upon binding to HilD, conformational changes are transmitted over the β-strands and affect the entire HilE domain. Hence, this HDX experiment on HilE does not allow us to draw further conclusions on the orientation or binding interface of HilE in the HilD-HilE heterodimer.

### Different Mechanisms of Regulating HilD Activity

HilE was previously shown to be dispensable for repression of HilD by *cis*-2-unsaturated LCFAs[33], as these compounds bind directly to HilD. We investigated whether competition exists between HilE and LCFAs for HilD, or if a possible additive effect exists between these negative regulators to repress HilD activity.

We first investigated the binding of HilD and HilE in the presence of oleic acid. Whilst the addition of DMSO had no effect on this interaction (Kd: 1.70 ± 0.50 μM), oleic acid prevented binding to HilE (Fig 7A). A reverse assay setup, probing oleic acid binding to the HilD/HilE complex, yielded an apparent K_d_ of 350 μM for oleic acid binding (Fig 7B). Changes in thermophoresis were only detected at oleic acid concentrations > 40 μM, significantly higher than the HilE concentration (10 μM), indicating that oleic acid is unable to bind to the HilD-HilE complex. Taken together, this shows that only one of these regulators can bind to, and regulate, HilD at a time and that the two regulatory mechanisms exist independently of one another.

**Figure 7.**
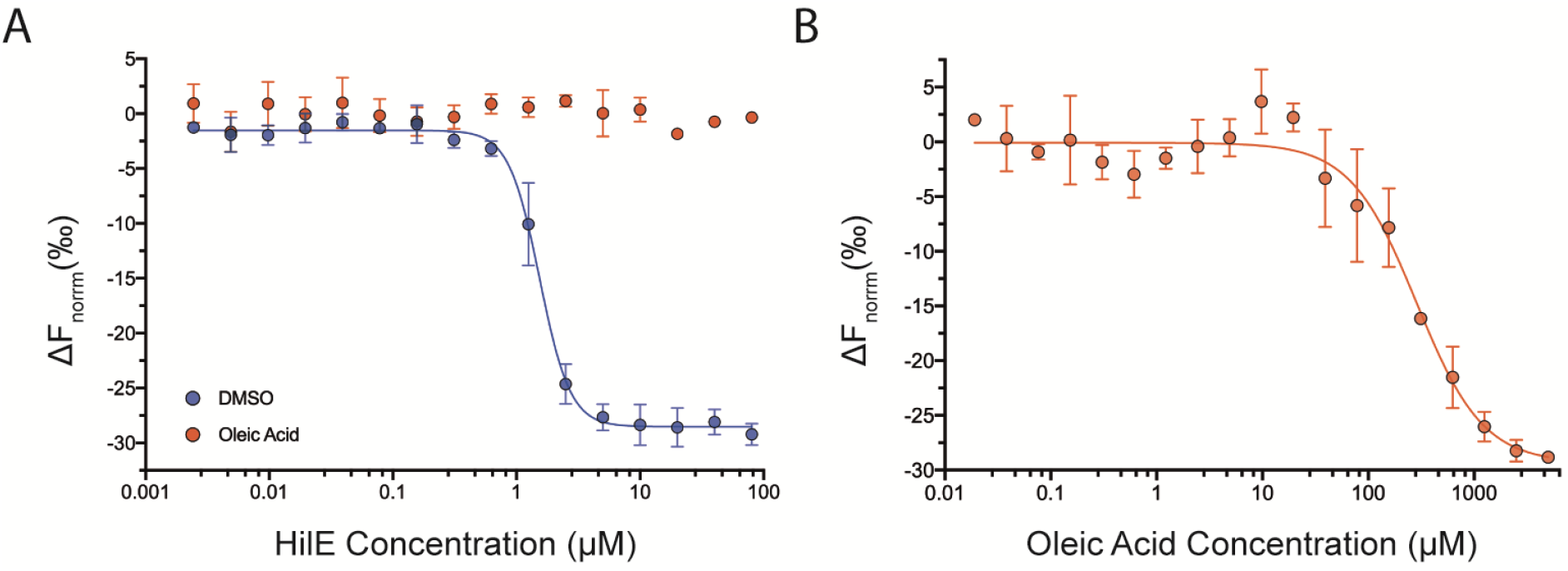
HilE and LCFAs bind independently to HilD. (**A**) MST dose-response curves for binding to HilE to HilD. GFP-HilD was first incubated with either 100 μM oleic acid (orange) or 1% DMSO (blue) and then increasing concentrations of HilE. (**B**) Oleic acid was titrated against a complex of HilD/HilE, constituted by mixing GFP-HilD (50 nM) with HilE (10 μM). An MST on-time of 5 seconds was used for analysis. Dose response curves were plotted using an MST on-time of 1.5 (A) or 5 (B) seconds. Error bars represent data obtained from three replicate experiments.

## Discussion

Long chain fatty acids regulate the function of several AraC/XylS transcription factors[40,41], and structures of ToxT and Rns show these ligands bind to a common pocket at the interface of the two protein domains[14,15]. Using HDX-MS, we showed that the fatty acid binding mode is also conserved in HilD. Our computational model predicted the interaction of fatty acids with residues K264 and R267 of HilD and, as seen for ToxT and Rns, shows that whilst the pocket is conserved between regulators, the specific binding residues vary. Specific interactions formed between the bound fatty acid and residues situated on both HilD domains may constrain HilD to a more stable, closed conformation and we found that binding of oleic acid increases the melting temperature of HilD. This supports the hypothesis that regulation of AraC/XylS transcription factors by fatty acids occurs via a common dynamic allosteric mechanism[35], inhibiting protein dimerisation and binding to target DNA. HilD can bind a range of fatty acids with a chain length of at least 16 carbon atoms. Fatty acid mimetics that meet the structural requirements for binding and present an opportunity for optimisation of increasingly potent inhibitors of *Salmonella* virulence.

The activity of AraC/XylS transcription regulators may also be modulated through protein-protein interactions, and a conserved family of AraC negative regulators is widespread amongst pathogenetic bacteria species in which virulence genes are regulated by AraC/XylS proteins[42]. The *Salmonella* negative regulator HilE is instead homologous to Hcp proteins. Our results show that unlike other characterised Hcp proteins, HilE exists predominately as a monomer, and the deletion of an extended sequence shown to be critical for the oligomerisation of other Hcp family proteins supports the hypothesis that HilE diverged from an ancestral structural Hcp protein required for virulence to a regulator of such virulence genes.

HilE forms a 1:1 complex with HilD, to inhibit HilD homodimerisation and prevent binding to DNA. Our results indicate that HilE interacts with the dimerisation helix of HilD, directly replacing one of the HilD monomers constituting the dimer pair. LCFA-binding to HilD is expected to result in conformational changes in the dimerisation helix, as reported for ToxT, restricting this helix to an orientation that is incompatible with the binding of HilE. This contrasts to the *P. aeruginosa* ExsD, which inhibits the dimerisation of the AraC/XylS regulator ExsA but does not bind directly to the ExsA dimer interface[43], and our results therefore highlight a previously unreported mechanism for the regulation of AraC/XylS proteins. The apparent novelty of this mechanism presents an attractive prospect for highly specific HilD binders. In addition to small molecules targeting the HilE-HilD complex, peptide-based inhibitors could be designed that mimic HilE binding to inhibit HilD dimerisation and activity.

Binding affinity measurements showed that the HilD/HilE interaction is of higher affinity than that calculated for the homodimerisation of HilD, supporting the hypothesis that HilD is bound to HilE under normal, non-invasive conditions. This repressive effect would only be overcome once HilD is expressed above the level of available HilE, by the action of positive regulators under conditions suitable for invasion (i.e. at the intestinal epithelium). Our results show that the mechanism of LCFA-repression of HilD is independent and mutually exclusive from that of HilE. HilD can only activate *hilA* expression when all conditions surpassing these repressive effects are met simultaneously, underlining the level of control over the expression of virulence genes to ensure efficiency of *Salmonella* pathogenesis.

## Materials and methods

### Cloning of protein constructs for expression

Genes encoding the desired proteins were inserted into either pET-21a(+) (*hilC, hilE*) or pET-24a(+) (*hilD*). The *hilD* construct contained an N-terminal His6-SUMO tag for expression, whilst *hilC* and *hilE* contained an N-terminal His6 tag followed by a TEV protease cleavage site. The *hilC* and *hilE* fusion genes were synthesised and purchased from Synbio Technologies, and the pET-SUMO-HilD plasmid was a gift from Marc Erhardt.

To clone the GFP-HilD fusion protein, the *gfp* gene was amplified and BamHI (GGATCC) and EheI (GGCGCC) restriction sites inserted for ligation into the SUMO HilD expression vector, into which the corresponding restriction sites were also introduced.

The His6-tagged HilD N-terminal domain construct was cloned using Round-the-Horn PCR, using the primer pairs HilD_NTD_fwd/rev and His-NTD_fwd/rev (Table 3) to remove the DNA-binding domain and SUMO fusion tag, respectively. This construct comprised HilD residues 7-206, with a His6 tag in place of the first six N-terminal residues of the HilD sequence.

**Table 1.** Affinity values for the binding of LCFAs to HilD.

| Lipid | Shorthand | $K_d$ (μM) | 68% CI (±) | Response Amplitude | No. repeats |
| --- | --- | --- | --- | --- | --- |
| Myristoleic Acid | 9Z-14:1 | 566.70 | 59.46 | 53.96 | 4 |
| Palmitoleic Acid | 9Z-16:1 | 58.53 | 4.86 | 41.48 | 4 |
| Oleic Acid | 9Z-18:1 | 48.06 | 5.27 | 41.24 | 3 |
| Gadoleic Acid | 9Z-20:1 | 26.68 | 3.13 | 31.10 | 4 |
| Erucic Acid | 13Z-22:1 | 14.71 | 2.19 | 32.60 | 3 |
| Nervonic Acid | 15Z-24:1 | 17.99 | 1.84 | 33.80 | 3 |
| Methyl-oleate | 9Z-18:1 | 738.54 | 276 | 5.61 | 3 |

K<sub>d</sub> values were calculated from changes in normalised fluorescence ( $\Delta F_{\text{norm}}$ ) at an MST on-time of 1.5 seconds with increasing ligand concentrations, from at least three replicate runs. Replicates were merged and the K<sub>d</sub> confidence interval (68% confidence) calculated using the MO.Affinity Analysis v2.3 software (NanoTemper Technologies GmbH).

**Table 2.**
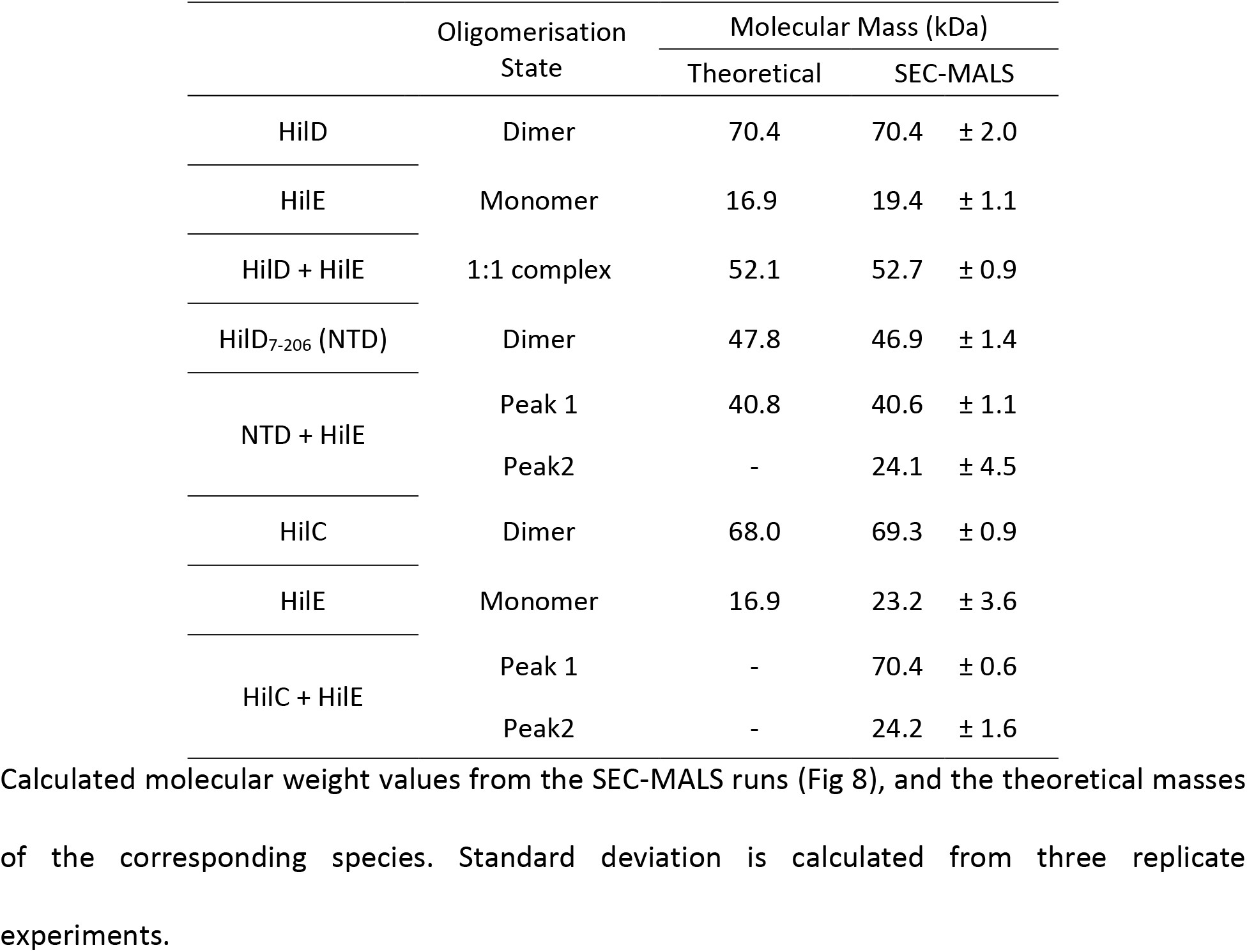
Molecular weight values determined from SEC-MALS.

**Table 3.**
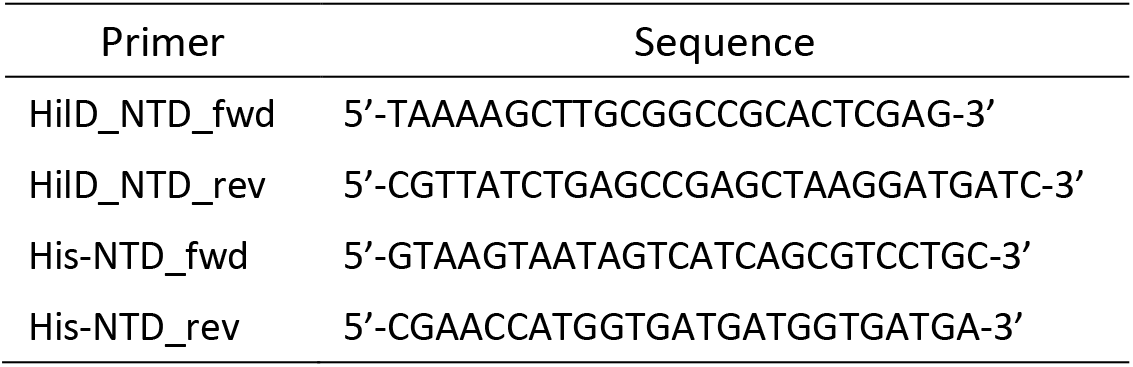
Primers used for Round-the-Horn PCR.

### Recombinant protein expression

Proteins were expressed in *E. coli* C41(DE3)[44] (or LEMO(DE3)[45] in the case of HilE) cells using lysogeny broth (LB) medium. An overnight culture was inoculated into LB medium and grown at 37 °C until an OD600nm of 0.6-0.8 was reached and induced by the addition of 0.5 mM isopropyl β-D-1-thiogalactopyranoside (IPTG). Cells were incubated with shaking overnight at 25 °C and collected by centrifugation (11,800 g, 4 °C).

### Protein purification

Pelleted cells were resuspended in a lysis buffer, supplemented with DNase and one cOmplete™ EDTA-free protease inhibitor cocktail tablet (Roche #11 873 580 001). Cells were lysed using a French press (2x, 1,000 psi) and centrifuged (95,000 g, 1 h, 4 °C). The resulting supernatant was filtered (0.40 μm) and loaded to a Ni-NTA column. The column was washed with 20% elution buffer, and the proteins eluted with an 100% elution buffer. The composition of buffers for protein purification is provided in Table 4.

**Table 4.**
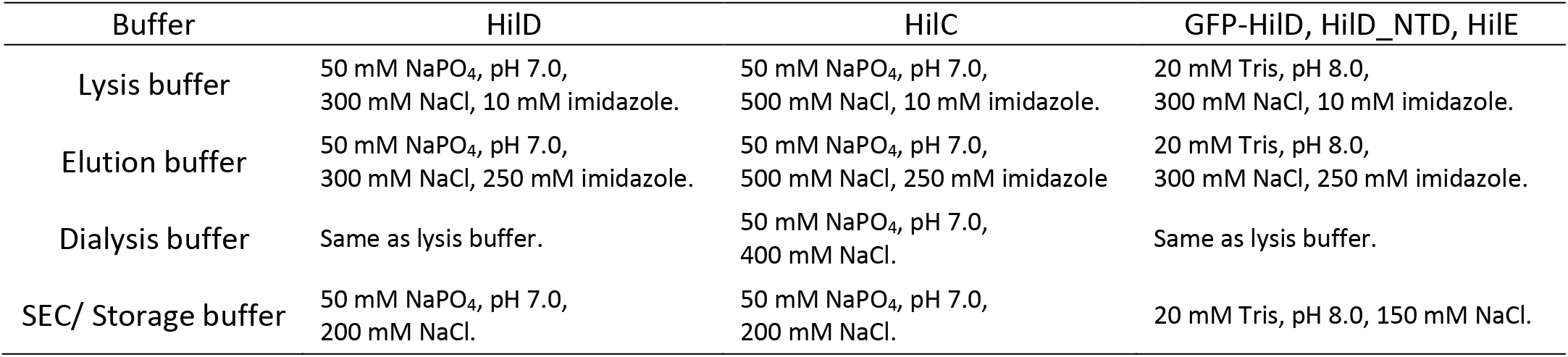
Buffers used for protein purification.

For HilD and GFP-HilD, the eluted proteins were supplemented with SUMO protease (600 μl, 0.4 mg ml^-1^) to cleave the His6-SUMO tag and dialyzed overnight at room temperature. The dialysed protein was reapplied to the Ni-NTA column, equilibrated with lysis buffer. The column was washed with 25% elution buffer to elute the cleaved protein. Proteins were then further purified by size-exclusion chromatography, using a HiLoad 26/60 Superdex 75 pg or HiLoad 26/60 Superdex 200 pg column for HilD and GFP-HilD, respectively, equilibrated with SEC buffer. Purified proteins were concentrated and stored in aliquots at −80 °C.

In the case of HilC and HilE, dialysis of proteins following the initial Ni-NTA column was performed overnight at 6 °C, with and the protein mixture supplemented with TEV protease (1 mg) prior to dialysis. The dialysed protein was reapplied to the Ni-NTA column as described for HilD, and column wash fractions containing the desired protein were combined and dialysed twice against storage buffer at 6 °C prior to storage.

For the HilD_NTD construct, following the initial Ni-NTA affinity purification step, the protein was concentrated and loaded to HiLoad 26/60 Superdex 75 pg column, equilibrated with SEC buffer.

### SEC-MALS

SEC-MALS experiments were performed using Superdex 75 Increase 10/300 GL column (Cytiva) coupled to a miniDAWN Tristar Laser photometer (Wyatt) and a RI-2031 differential refractometer (JASCO). 50 μl samples of 100 μM protein (50 μM in the case of HilC) were loaded to the SEC column, equilibrated with SEC buffer (50 mM NaPO4 pH 7.0, 200 mM NaCl), and separated using a flow rate of 0.5 ml min^-1^. Data analysis was carried out with ASTRA v7.3.0.18 software (Wyatt).

### NanoDSF

Protein samples were heated from 20-95 °C, with a temperature gradient of 0.5 °C min^-1^. Melting temperatures were calculated from changes in the fluorescence ratio (350/330 nm), using PR.Stability Analysis v1.0.3 software (NanoTemper Technologies). To assess ligand-induced effects on HilD stability, oleic acid was diluted 1:100 (final concentration 50 μM) into 20 μM HilD (in SEC buffer) to give a final DMSO concentration of 1% (v/v), and incubated for 20 min at room temperature.

### MST

All MST measurements were performed on a NanoTemper Monolith NT.115 with a Nano BLUE/RED Detector using MO.Control v1.6. MST runs were performed at 25 °C, with an excitation power of 60% and MST power set to medium. Data were analysed using the MO.Affinity Analysis v2.3 software, and affinity constants were calculated using the K_d_ model.

MST experiments were performed with 50 nM GFP-HilD and increasing concentrations of the respective ligand in either Tris MST assay buffer (20 mM Tris, pH 8.0, 150 mM NaCl, 0.1% (v/v) Pluronic) for HilE/ oleic acid binding, or NaPO4 MST assay buffer (50 mM NaPO4, pH 7.0, 200 mM NaCl, 0.1% (v/v) Pluronic) for HilD dimerisation runs. For HilD dimerisation and HilE binding, a two-fold serial dilution of proteins was performed in the corresponding protein storage buffer. Dilution series of fatty acids was prepared in ethanol, and subsequently diluted 1:50 into assay buffer. All ligand samples were then mixed 1:1 with 100 nM GFP-HilD (diluted in assay buffer), incubated together for 10 min at room temperature, centrifuged for 5 min and loaded to standard capillaries (Nanotemper, #MO-K022). To investigate competitive effects of different ligands, GFP-HilD was pre-incubated with the competing ligand for ≥ 10 min at room temperature, before mixing with the second ligand (of varying concentration).

### Computational methods

#### Protein structure prediction

The structural model of HilD (UniProt ID: P0CL08) was retrieved from the AlphaFold Protein Structure Database[46]. The structure of HilE was predicted using the following publicly-available prediction structure prediction servers: Alphafold2[47], RoseTTAFold[48] and tFold (https://drug.ai.tencent.com). Complex structures of the HilD homodimer and HilD-HilE heterodimer were predicted using AlphaFold-Multimer (v2.2.0)[31]. All structure models can be found in the supplementary material.

#### Binding site prediction and molecular docking

System preparation and docking calculations were performed using the Schrödinger Drug Discovery suite for molecular modelling (version 2022.1). Protein-ligand complex was prepared with the Protein Preparation Wizard to fix protonation states of amino acids, add hydrogens, and fix missing side-chain atoms. All ligands for docking were drawn using Maestro and prepared using LigPrep[49] to generate the 3D conformation, adjust the protonation state to physiological pH (7.4), and calculate the partial atomic charges with the OPLS4 force field. Docking studies with the prepared ligands were performed using Glide (Glide V7.7)[50,51] with the flexible modality of induced-fit docking with extra precision (XP), followed by a side-chain minimization step using Prime. Ligands were docked within a grid around 12 Å from the centroid of the predicted binding site pocket, determined using SiteMap.

#### HDX

Two different HDX experiments were conducted on HilD (datasets 1 and 2 in Dataset S4, respectively) To investigate the impact of oleic acid on HilD conformation (dataset 1), HilD was supplemented with 1% (v/v) of either DMSO or oleic acid (10 mM in DMSO) yielding a final concentration of 100 μM oleic acid in the sample. For experiments probing the HilD/HilE interaction (dataset 2), samples contained either individual HilD or HilE, or the HilD/HilE complex (all proteins at 25 μM final concentration), which was established prior to HDX-MS by purification using a Superdex 75 Increase 10/300 GL column (Cytiva) equilibrated in SEC buffer (50 mM NaH2PO4/Na2HPO4 pH 7.0, 200 mM NaCl). Final assay concentrations of HilD and/or HilE in all experiments were 25 μM. All samples were stored in a cooled tray (1 °C) until measurement.

Preparation of the HDX reactions was aided by a two-arm robotic autosampler (LEAP technologies). 7.5 μl of protein sample (see above) was mixed with 67.5 μl of SEC buffer prepared with 99.9% D2O to initiate the hydrogen exchange reaction. After incubation at 25 °C for 10, 30, 100, 1,000 or 10,000 seconds, 55 μl of the HDX reaction was withdrawn and added to 55 μl of pre-dispensed quench buffer (400 mM KH2PO4/H3PO4, pH 2.2, 2 M guanidine-HCl) kept at 1 °C. 95 μl of the resulting mixture was injected into an ACQUITY UPLC M-Class System with HDX Technology (Waters)[52]. Undeuterated protein samples were prepared similarly (incubation for approximately 10 s at 25 °C) through 10-fold dilution of protein samples with H_2_O-containing SEC buffer. The injected samples were flushed out of the loop (50 μl) with H_2_O + 0.1% (v/v) formic acid (100 μl min^-1^) and guided to a protease column (2 mm x 2 cm) containing proteases immobilized to the bead material, which was kept at 12 °C. For each protein state and timepoint, three replicates (individual HDX reactions) were digested with porcine pepsin, while another three replicates were digested with a column filled with a 1:1 mixture of protease type XVIII from *Rhizopus spp.* and protease type XIII from *Aspergillus saitoi.* In both cases, the resulting peptides were trapped on an AQUITY UPLC BEH C18 1.7 μm 2.1 x 5 mm VanGuard Pre-column (Waters) kept at 0.5 °C. After 3 min of digestion and trapping, the trap column was placed in line with an ACQUITY UPLC BEH C18 1.7 μm 1.0 x 100 mm column (Waters), and the peptides eluted at 0.5 °C using a gradient of buffers A (H_2_O + 0.1% (v/v) formic acid) and B (acetonitrile + 0.1% (v/v) formic acid) at a flow rate of 60 μl min^-1^ as follows: 0-7 min: 95-65% A; 7-8 min: 65-15% A; 8-10 min: 15% A; 10-11 min: 5% A; 11-16 min: 95% A. The eluted proteins were guided to a G2-Si HDMS mass spectrometer with ion mobility separation (Waters), and peptides ionized with an electrospray ionization source (250 °C capillary temperature, spray voltage 3.0 kV) and mass spectra acquired in positive ion mode over a range of 50 to 2,000 m/z in enhanced high definition MS (HDMS^E^) or high definition MS (HDMS) mode for undeuterated and deuterated samples, respectively[53,54]. [Glu1]-Fibrinopeptide B standard (Waters) was employed for lock-mass correction. During separation of the peptide mixtures on the ACQUITY UPLC BEH C18 column, the protease column was washed three times with 80 μl of wash solution (0.5 M guanidine hydrochloride in 4% (v/v) acetonitrile,) and blank injections performed between each sample to reduce peptide carry-over.

Peptide identification and analysis of deuterium incorporation were carried out with ProteinLynx Global SERVER (PLGS, Waters) and DynamX 3.0 softwares (Waters) as described previously[55]. In summary, peptides were identified with PLGS from the undeuterated samples acquired with HDMS^E^ by employing low energy, elevated energy, and intensity thresholds of 300, 100 and 1,000 counts, respectively. Identified ions were matched to peptides with a database containing the amino acid sequence of HilD, HilE, porcine pepsin, and their reversed sequences with the following search parameters: peptide tolerance = automatic; fragment tolerance = automatic; min fragment ion matches per peptide = 1; min fragment ion matches per protein = 7; min peptide matches per protein = 3; maximum hits to return = 20; maximum protein mass = 250,000; primary digest reagent = non-specific; missed cleavages = 0; false discovery rate = 100. Only peptides that were identified in all undeuterated samples and with a minimum intensity of 30,000 counts, a maximum length of 30 amino acids, a minimum number of three products with at least 0.1 product per amino acid, a maximum mass error of 25 ppm and retention time tolerance of 0.5 minutes were considered for further analysis. Deuterium incorporation into peptides was quantified with DynamX 3.0 software (Waters). Hereby, the datasets generated with pepsin digestion or after digestions with proteases type XIII and XVIII were pooled. All spectra were manually inspected and, if necessary, peptides omitted (e.g., in case of low signal-to-noise ratio or presence of overlapping peptides).

The observable maximal deuterium uptake of a peptide (see Dataset S4) was calculated by the number of residues minus one (for the N-terminal residue) minus the number of proline residues contained in the peptide. For the calculation of HDX in per cent the absolute HDX was divided by the theoretical maximal deuterium uptake multiplied by 100. To render the residue specific HDX differences from overlapping peptides for any given residue of HilD or HilE, the shortest peptide covering this residue was employed. Where multiple peptides were of the shortest length, the peptide with the residue closest to the peptide’s C-terminus was utilized.

#### EMSA

EMSAs were performed using a 62 base pair dsDNA fragment of the *hilA* promoter encompassing the A1 binding site[56]. Double stranded DNA fragments were generated by boiling complementary primers together at 95°C for 10 min, before slowly cooling to room temperature. The forward primer was modified with a 5’-Cy5 fluorescent dye for detection. 50 nM of labelled DNA was incubated with increasing concentrations of protein in EMSA buffer (20 mM Tris, pH 8.0, 100 mM KCl, 100 μM EDTA, 3% glycerol). Samples were incubated at 37°C for 15 min, supplemented with diluted DNA loading dye, and separated on a 1.5 mm thick, 6% TBE gel at 6°C at a constant voltage of 100 V. Gels were imaged using a ChemiDoc™MP imaging system (Bio-Rad Inc). Primer sequences were as follows: Fwd: ([Cyanine5]GGGAGTAAAGAAAAGACGATATCATTATTTTGCAAAAAAATATAAAAATAAGCGCACCATTA), Rev: (TAATGGTGCGCTTATTTTTATATTTTTTTGCAAAATAATGATATCGTCTTTTCTTTACTCCC).

## Contributions

J.D.J. and M.D.H. conceptualized the study. J.D.J. performed cloning, protein expression and purification, protein assays, and analysed collected data. W.S. performed HDX-MS experiments and analysed the data. T.K. performed computational docking simulations. G.B., A.P., S.W., M.D.H. were responsible for supervision, project administration and funding acquisition. J.D.J. wrote the paper, with input from all other authors.

## Acknowledgments

We thank Marc Erhardt for the pET-SUMO-HilD plasmid, Iwan Grin for help with optimising HilD expression and purification and Abdelhakim Boudrioua for scientific feedback and discussions. We are also grateful to Reinhard Albrecht and Nick Mozer for support in protein purification, and Hadeer Elhabashy for assistance in modelling of the HilD/HilE complex. The authors would like to thank CSC-Finland for the generous computational resources. Structure models were computed using the BMBF-funded de.NBI Cloud within the German Network for Bioinformatics Infrastructure (de.NBI) (031A532B, 031A533A, 031A533B, 031A534A, 031A535A, 031A537A, 031A537B, 031A537C, 031A537D, 031A538A). Figure 1 and cartoons in figure 2D were created with Biorender.com.

This work was supported by the Baden-Württemberg Foundation grant BWST_WSF-018 (to A.P., S.W. and M.D.H.) and institutional funds of the Max Planck Society. T.K. is financed by the iFIT (EXC2180 – 390900677), Fortüne initiative (NR.2613-0), the cluster of excellence EXC2124 Controlling Microbes to Fight Infections (CMFI), and the Federal Ministry of Education and Research (BMBF) and the Baden-Württemberg Ministry of Science as part of the Excellence Strategy of the German Federal and State Governments – Germany, by the means of the program TüCAD2. We acknowledge support by the German Research Council (DFG) through the core facility for HDX-MS (project 324652314 to G.B.).

## Supporting information

**Figure S1.**
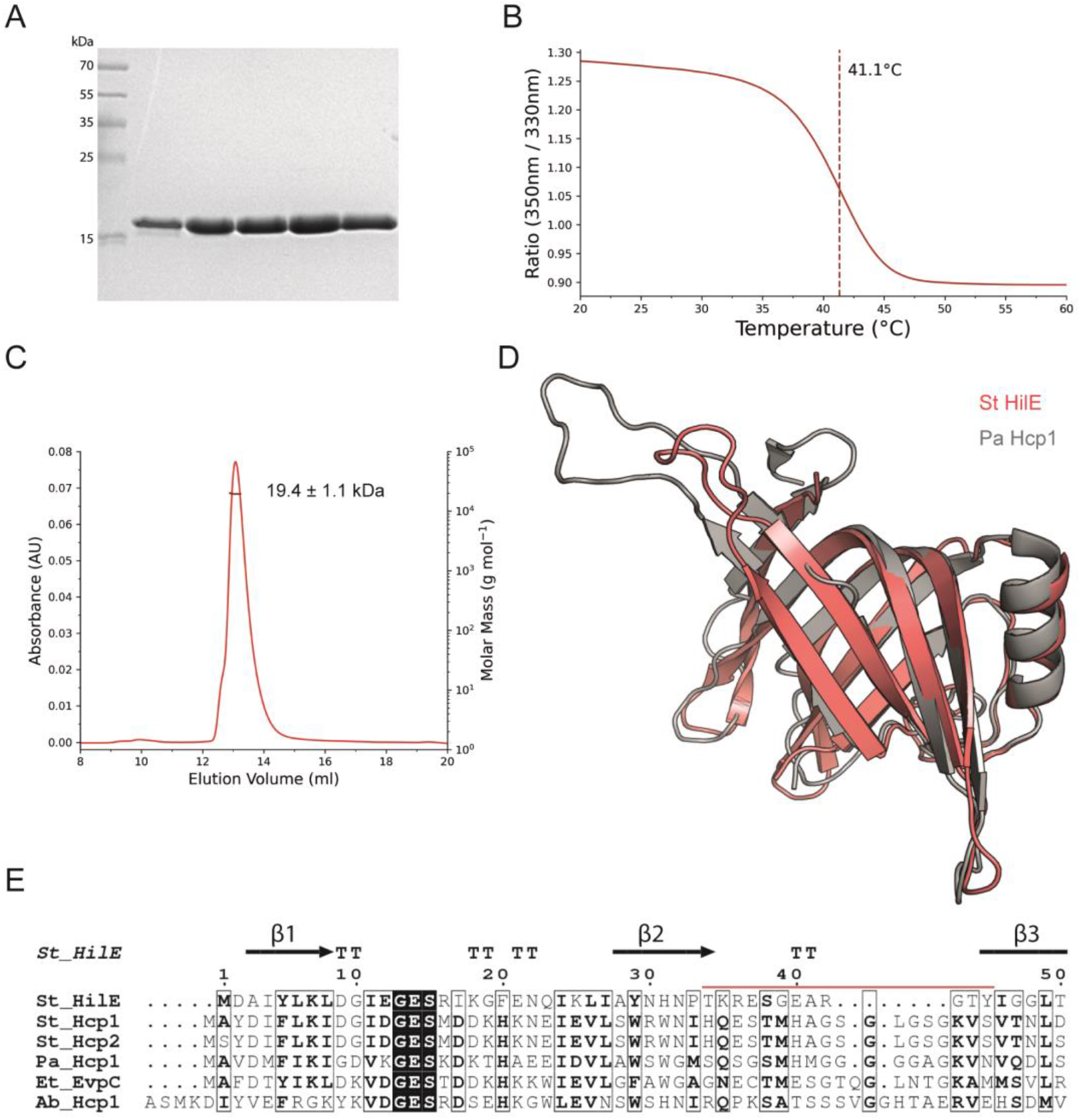
HilE exists exclusively as a monomer in solution. (**A**) Coomassie-stained SDS-PAGE gel for fractions eluted from the final purification step of HilE. (**B**) NanoDSF unfolding profile of HilE (1 mg ml^-1^). (**C**) SEC-MALS elution profile of HilE. HilE (100 μM) was loaded to a S75 10/300 increase column, and absorbance was constantly monitored at 280 nm. Molecular mass of the eluted protein was calculated from the static light scattering. (**D**) Structural alignment of HilE (tfold prediction, red) with *Pseudomonas aeruginos* Hcp1 (PDB: 1Y12; grey). (**E**) Multiple sequence alignment of Hcp proteins, identified using HHPred, was performed using ClustalΩ and highlights the shortened length of the loop in HilE (residues constituting the loop in HilE are marked by the red line). β-strands, as predicted in the structure of HilE, are denoted by the arrows above the alignment.

**Figure S2.**
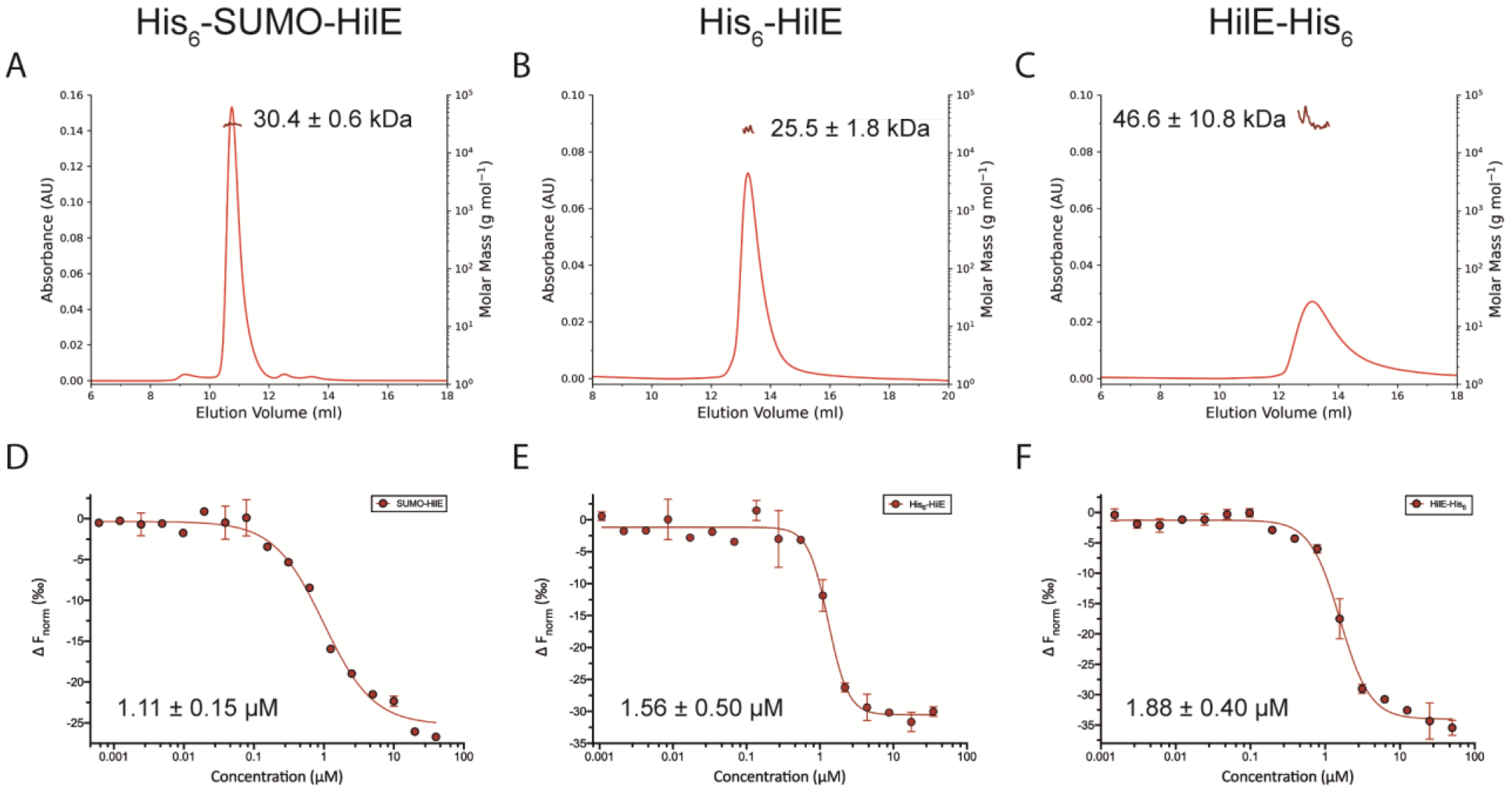
Oligomerisation and activity of additional purified HilE constructs. (**A-C**) SEC-MALS elution profiles of the three purified HilE constructs. Molecular weight values from light scattering measurements were calculated from three replicate runs for each construct. (**D-F**) Corresponding dose-response binding curves for the binding of each of the HilE constructs to GFP-HilD in an MST assay. K_d_ affinity values were calculated from two repeat measurements, and an MST on-time of 2.5 seconds used for analysis.

**Figure S3.**
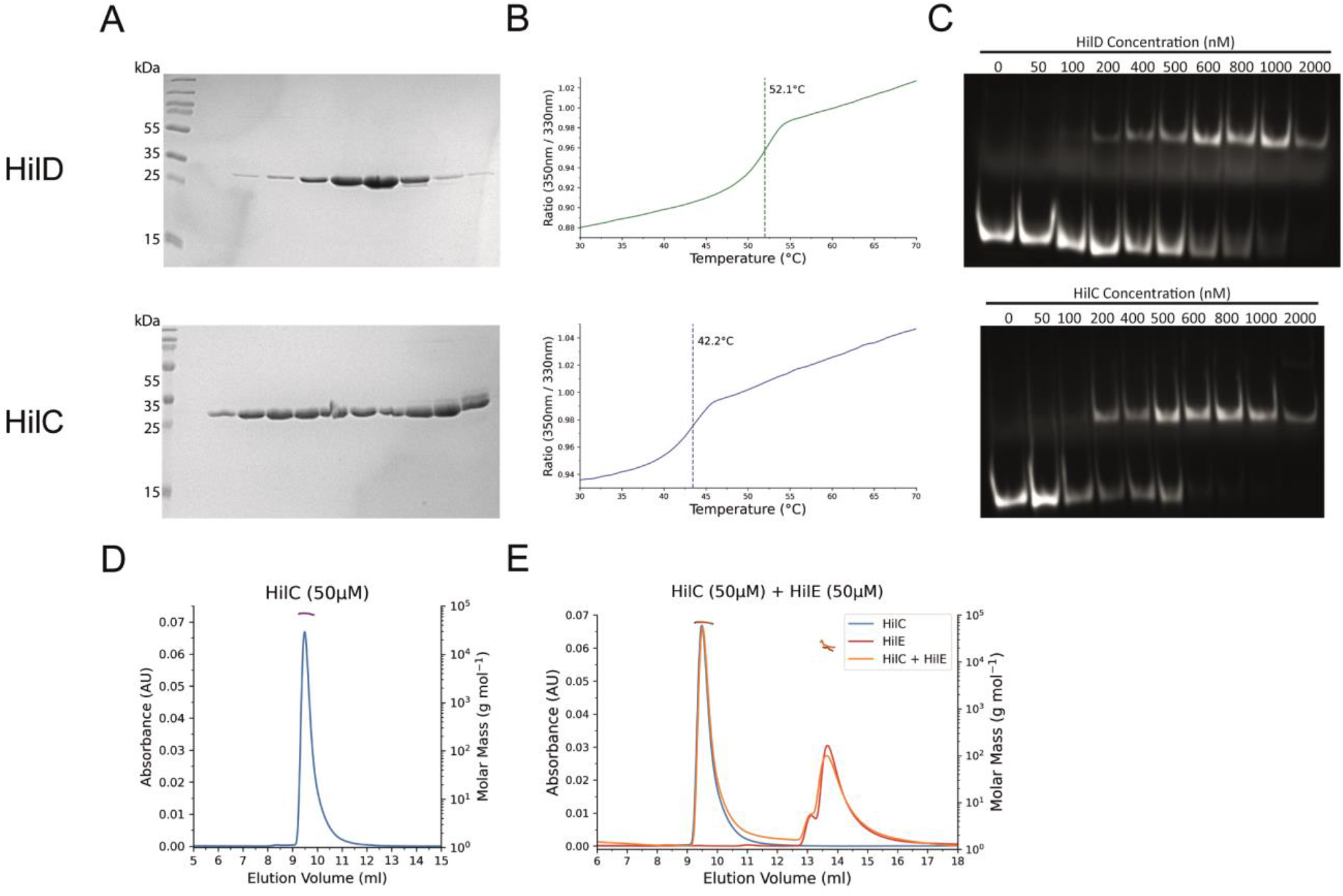
Biophysical characterisation of HilD and HilC. (**A**) Coomassie-stained SDS-PAGE gel for fractions eluted from the final purification step of HilD (top) and HilC (bottom). (**B**) NanoDSF unfolding profile of HilD (green, top) and HilC (blue, bottom). A concentration of 20 μM was used for both proteins. (**C**) EMSA showing binding of HilD (top) and HilC (bottom) to the *hilA* promoter. All lanes contain 50 nM of a DNA fragment encompassing the A1 HilD binding site within the promoter and increasing protein concentrations, as indicated. DNA is labelled with a Cy5 fluorophore at the 5’ end of the forward strand for image detection. (**D-E**) SEC-MALS profiles of (D) HilC and (E) HilC/HilE. Protein concentrations of 50 μM were used for all samples, due to the lower solubility and observed precipitation of HilC at higher concentrations. Molecular weight values were calculated from three repeat experiments and are displayed in Table 2.

**Dataset S4. Overview HDX-MS conditions and deuterium incorporation of peptides.**

**Dataset S5. Predicted AlphaFold Multimer structures of the HilD-HilD homodimer.**

**Dataset S6. Predicted AlphaFold Multimer structures of the HilD-HilE heterodimer.**

## References

1. Fàbrega A, Vila J. Salmonella enterica serovar Typhimurium skills to succeed in the host: Virulence and regulation. Clin Microbiol Rev. 2013;26: 308–341. doi:10.1128/CMR.00066-12

2. Ilyas B, Tsai CN, Coombes BK. Evolution of Salmonella-host cell interactions through a dynamic bacterial genome. Front Cell Infect Microbiol. 2017;7. doi:10.3389/fcimb.2017.00428

3. Lerminiaux NA, MacKenzie KD, Cameron ADS. Salmonella Pathogenicity Island 1 (SPI-1): The Evolution and Stabilization of a Core Genomic Type Three Secretion System. Microorganisms. 2020;8. doi:10.3390/microorganisms8040576

4. Haraga A, Ohlson MB, Miller SI. Salmonellae interplay with host cells. Nat Rev Microbiol. 2008;6: 53–66. doi:10.1038/nrmicro1788

5. Hansen-Wester I, Hensel M. Salmonella pathogenicity islands encoding type III secretion systems. Microbes and Infection. 2001. doi:10.1016/S1286-4579(01)01411-3

6. Bajaj V, Hwang C, Lee CA. hilA is a novel ompR/toxR family member that activates the expression of Salmonella typhimurium invasion genes. Mol Microbiol. 1995;18: 715–27. doi:10.1111/j.1365-2958.1995.mmi_18040715.x

7. Eichelberg K, Galán JE. Differential regulation of Salmonella typhimurium type III secreted proteins by pathogenicity island 1 (SPI-1)-encoded transcriptional activators InvF and hilA. Infect Immun. 1999;67: 4099–105. doi:10.1128/IAI.67.8.4099-4105.1999

8. Lostroh CP, Lee CA. The Salmonella pathogenicity island-1 type III secretion system. Microbes and Infection. 2001. pp. 1281–1291. doi:10.1016/S1286-4579(01)01488-5

9. Olekhnovich IN, Kadner RJ. DNA-binding activities of the HilC and HilD virulence regulatory proteins of Salmonella enterica serovar Typhimurium. J Bacteriol. 2002;184: 4148–4160. doi:10.1128/JB.184.15.4148-4160.2002

10. Olekhnovich IN, Kadner RJ. Role of nucleoid-associated proteins Hha and H-NS in expression of Salmonella enterica activators HilD, HilC, and RtsA required for cell invasion. J Bacteriol. 2007;189: 6882–6890. doi:10.1128/JB.00905-07

11. Gallegos MT, Schleif R, Bairoch A, Hofmann K, Ramos JL. Arac/XylS family of transcriptional regulators. Microbiol Mol Biol Rev. 1997;61: 393–410. doi:10.1128/.61.4.393-410.1997

12. Perez-Rueda E, Hernandez-Guerrero R, Martinez-Nuñez MA, Armenta-Medina D, Sanchez I, Ibarra JA. Abundance, diversity and domain architecture variability in prokaryotic DNA-binding transcription factors. PLoS One. 2018;13: e0195332. doi:10.1371/journal.pone.0195332

13. Cortés-Avalos D, Martínez-Pérez N, Ortiz-Moncada MA, Juárez-González A, Baños-Vargas AA, Estrada-de los Santos P, et al. An update of the unceasingly growing and diverse AraC/XylS family of transcriptional activators. FEMS Microbiol Rev. 2021;45. doi:10.1093/femsre/fuab020

14. Lowden MJ, Skorupski K, Pellegrini M, Chiorazzo MG, Taylor RK, Kull FJ. Structure of Vibrio cholerae ToxT reveals a mechanism for fatty acid regulation of virulence genes. Proc Natl Acad Sci U S A. 2010;107: 2860–2865. doi:10.1073/pnas.0915021107

15. Midgett CR, Talbot KM, Day JL, Munson GP, Kull FJ. Structure of the master regulator Rns reveals an inhibitor of enterotoxigenic Escherichia coli virulence regulons. Sci Rep. 2021;11: 1–13. doi:10.1038/s41598-021-95123-2

16. Ellermeier CD, Ellermeier JR, Slauch JM. HilD, HilC and RtsA constitute a feed forward loop that controls expression of the SPI1 type three secretion system regulator hilA in Salmonella enterica serovar Typhimurium. Mol Microbiol. 2005;57: 691–705. doi:10.1111/j.1365-2958.2005.04737.x

17. Saini S, Ellermeier JR, Slauch JM, Rao C V. The role of coupled positive feedback in the expression of the SPI1 type three secretion system in Salmonella. PLoS Pathog. 2010;6: 1–16. doi:10.1371/journal.ppat.1001025

18. Golubeva YA, Sadik AY, Ellermeier JR, Slauch JM. Integrating global regulatory input into the Salmonella pathogenicity Island 1 type III secretion system. Genetics. 2012;190: 79–90. doi:10.1534/genetics.111.132779

19. Golubeva YA, Ellermeier JR, Chubiz JEC, Slauch JM. Intestinal long-chain fatty acids act as a direct signal to modulate expression of the Salmonella pathogenicity island 1 type III secretion system. MBio. 2016;7: 1–9. doi:10.1128/mBio.02170-15

20. Chowdhury R, Bitar PDP, Keresztes I, Condo AM, Altier C. A diffusible signal factor of the intestine dictates Salmonella invasion through its direct control of the virulence activator HilD. PLoS Pathog. 2021;17: 1–20. doi:10.1371/journal.ppat.1009357

21. Chowdhury R, Pavinski Bitar PD, Adams MC, Chappie JS, Altier C. AraC-type regulators HilC and RtsA are directly controlled by an intestinal fatty acid to regulate Salmonella invasion. Mol Microbiol. 2021;116: 1464–1475. doi:10.1111/mmi.14835

22. Yang X, Stein KR, Hang HC. Anti-infective bile acids bind and inactivate a Salmonella virulence regulator. Nat Chem Biol. 2022. doi:10.1038/s41589-022-01122-3

23. Wu Y, Yang X, Zhang D, Lu C. Myricanol Inhibits the Type III Secretion System of Salmonella enterica Serovar Typhimurium by Interfering With the DNA-Binding Activity of HilD. Front Microbiol. 2020;11: 1–11. doi:10.3389/fmicb.2020.571217

24. Shi Y, Chen X, Shu J, Liu Y, Zhang Y, Lv Q, et al. Harmine, an inhibitor of the type III secretion system of Salmonella enterica serovar Typhimurium. Front Cell Infect Microbiol. 2022;12. doi:10.3389/fcimb.2022.967149

25. Baxter MA, Fahlen TF, Wilson RL, Jones BD. HilE interacts with HilD and negatively regulates hilA transcription and expression of the Salmonella enterica serovar Typhimurium invasive phenotype. Infect Immun. 2003;71: 1295–1305.

26. Grenz JR, Chubiz JEC, Thaprawat P, Slauch JM. HilE regulates HilD by blocking DNA binding in Salmonella enterica serovar Typhimurium. J Bacteriol. 2018;200: 1–13. doi:10.1128/JB.00750-17

27. Paredes-Amaya CC, Valdés-García G, Juárez-González VR, Rudiño-Piñera E, Bustamante VH. The Hcp-like protein HilE inhibits homodimerization and DNA binding of the virulence-associated transcriptional regulator HilD in Salmonella. J Biol Chem. 2018;293: 6578–6592. doi:10.1074/jbc.RA117.001421

28. Egan SM, Schleif RF. DNA-dependent renaturation of an insoluble DNA binding protein. Identification of the RhaS binding site at rhaBAD. J Mol Biol. 1994;243: 821–9. doi:10.1006/jmbi.1994.1684

29. Schäper S, Steinchen W, Krol E, Altegoer F, Skotnicka D, Søgaard-Andersen L, et al. AraC-like transcriptional activator CuxR binds c-di-GMP by a PilZ-like mechanism to regulate extracellular polysaccharide production. Proc Natl Acad Sci U S A. 2017;114: E4822–E4831. doi:10.1073/pnas.1702435114

30. Narm KE, Kalafatis M, Slauch JM. HilD, HilC, and RtsA form homodimers and heterodimers to regulate expression of the salmonella pathogenicity island i type iii secretion system. J Bacteriol. 2020;202: 1–13. doi:10.1128/JB.00012-20

31. Evans R, O’Neill M, Pritzel A, Antropova N, Senior A, Green T, et al. Protein complex prediction with AlphaFold-Multimer. bioRxiv. 2021; 2021.10.04.463034. doi:10.1101/2021.10.04.463034

32. Shrestha M, Xiao Y, Robinson H, Schubot FD. Structural analysis of the regulatory domain of ExsA, a key transcriptional regulator of the type three secretion system in Pseudomonas aeruginosa. PLoS One. 2015;10: 1–17. doi:10.1371/journal.pone.0136533

33. Bosire EM, Eade CR, Schiltz CJ, Reid AJ, Troutman J, Chappie JS, et al. Diffusible Signal Factors Act through AraC-Type Transcriptional Regulators as Chemical Cues To Repress Virulence of Enteric Pathogens. Infect Immun. 2020;88: 1–14. doi:10.1128/IAI.00226-20

34. Shakhnovich EA, Hung DT, Pierson E, Lee K, Mekalanos JJ. Virstatin inhibits dimerization of the transcriptional activator ToxT. Proc Natl Acad Sci U S A. 2007;104: 2372–2377. doi:10.1073/pnas.0611643104

35. Cruite JT, Kovacikova G, Clark KA, Woodbrey AK, Skorupski K, Kull FJ. Structural basis for virulence regulation in Vibrio cholerae by unsaturated fatty acid components of bile. Commun Biol. 2019;2: 1–9. doi:10.1038/s42003-019-0686-x

36. Childers BM, Cao X, Weber GG, Demeler B, Hart PJ, Klose KE. N-terminal Residues of the Vibrio cholerae Virulence Regulatory Protein ToxT Involved in Dimerization and Modulation by Fatty Acids. J Biol Chem. 2011;286: 28644–28655. doi:10.1074/jbc.M111.258780

37. Lim YT, Jobichen C, Wong J, Limmathurotsakul D, Li S, Chen Y, et al. Extended Loop Region of Hcp1 is Critical for the Assembly and Function of Type VI Secretion System in Burkholderia pseudomallei. Sci Rep. 2015;5: 8235. doi:10.1038/srep08235

38. Lin Q-P, Gao Z-Q, Geng Z, Zhang H, Dong Y-H. Crystal structure of the putative cytoplasmic protein STM0279 (Hcp2) from *Salmonella typhimurium*. Acta Crystallogr Sect F Struct Biol Commun. 2017;73: 463–468. doi:10.1107/S2053230X17010512

39. Krissinel E, Henrick K. Inference of Macromolecular Assemblies from Crystalline State. J Mol Biol. 2007;372: 774–797. doi:10.1016/j.jmb.2007.05.022

40. Chatterjee A, Dutta PK, Chowdhury R. Effect of Fatty Acids and Cholesterol Present in Bile on Expression of Virulence Factors and Motility of *Vibrio cholerae*. Infect Immun. 2007;75: 1946–1953. doi:10.1128/IAI.01435-06

41. Day J, Kovacikova G, Taylor RK, Kull FJ. Unsaturated Fatty Acid Regulation of AraC/XylS Transcription Factors. Biophys J. 2014;106: 497a. doi:10.1016/j.bpj.2013.11.2779

42. Santiago AE, Yan MB, Tran M, Wright N, Luzader DH, Kendall MM, et al. A large family of anti-activators accompanying XylS/AraC family regulatory proteins. Mol Microbiol. 2016;101: 314–332. doi:10.1111/mmi.13392

43. Shrestha M, Bernhards RC, Fu Y, Ryan K, Schubot FD. Backbone Interactions Between Transcriptional Activator ExsA and Anti-Activator ExsD Facilitate Regulation of the Type III Secretion System in Pseudomonas aeruginosa. Sci Rep. 2020;10: 9881. doi:10.1038/s41598-020-66555-z

44. Miroux B, Walker JE. Over-production of Proteins inEscherichia coli: Mutant Hosts that Allow Synthesis of some Membrane Proteins and Globular Proteins at High Levels. J Mol Biol. 1996;260: 289–298. doi:10.1006/jmbi.1996.0399

45. Wagner S, Klepsch MM, Schlegel S, Appel A, Draheim R, Tarry M, et al. Tuning *Escherichia coli* for membrane protein overexpression. Proc Natl Acad Sci. 2008;105: 14371–14376. doi:10.1073/pnas.0804090105

46. Varadi M, Anyango S, Deshpande M, Nair S, Natassia C, Yordanova G, et al. AlphaFold Protein Structure Database: massively expanding the structural coverage of protein-sequence space with high-accuracy models. Nucleic Acids Res. 2022;50: D439–D444. doi:10.1093/nar/gkab1061

47. Jumper J, Evans R, Pritzel A, Green T, Figurnov M, Ronneberger O, et al. Highly accurate protein structure prediction with AlphaFold. Nature. 2021;596: 583–589. doi:10.1038/s41586-021-03819-2

48. Baek M, DiMaio F, Anishchenko I, Dauparas J, Ovchinnikov S, Lee GR, et al. Accurate prediction of protein structures and interactions using a three-track neural network. Science (80-). 2021;373: 871–876. doi:10.1126/science.abj8754

49. Shelley JC, Cholleti A, Frye LL, Greenwood JR, Timlin MR, Uchimaya M. Epik: a software program for pK a prediction and protonation state generation for drug-like molecules. J Comput Aided Mol Des. 2007;21: 681–691. doi:10.1007/s10822-007-9133-z

50. Friesner RA, Banks JL, Murphy RB, Halgren TA, Klicic JJ, Mainz DT, et al. Glide: A New Approach for Rapid, Accurate Docking and Scoring. 1. Method and Assessment of Docking Accuracy. J Med Chem. 2004;47: 1739–1749. doi:10.1021/jm0306430

51. Friesner RA, Murphy RB, Repasky MP, Frye LL, Greenwood JR, Halgren TA, et al. Extra Precision Glide: Docking and Scoring Incorporating a Model of Hydrophobic Enclosure for Protein-Ligand Complexes. J Med Chem. 2006;49: 6177–6196. doi:10.1021/jm051256o

52. Wales TE, Fadgen KE, Gerhardt GC, Engen JR. High-speed and high-resolution UPLC separation at zero degrees Celsius. Anal Chem. 2008;80: 6815–20. doi:10.1021/ac8008862

53. Geromanos SJ, Vissers JPC, Silva JC, Dorschel CA, Li G-Z, Gorenstein M V., et al. The detection, correlation, and comparison of peptide precursor and product ions from data independent LC-MS with data dependant LC-MS/MS. Proteomics. 2009;9: 1683–1695. doi:10.1002/pmic.200800562

54. Li G-Z, Vissers JPC, Silva JC, Golick D, Gorenstein M V., Geromanos SJ. Database searching and accounting of multiplexed precursor and product ion spectra from the data independent analysis of simple and complex peptide mixtures. Proteomics. 2009;9: 1696–1719. doi:10.1002/pmic.200800564

55. Osorio-Valeriano M, Altegoer F, Steinchen W, Urban S, Liu Y, Bange G, et al. ParB-type DNA Segregation Proteins Are CTP-Dependent Molecular Switches. Cell. 2019;179: 1512–1524.e15. doi:10.1016/j.cell.2019.11.015

56. Olekhnovich IN, Kadner RJ. Crucial roles of both flanking sequences in silencing of the hilA promoter in Salmonella enterica. J Mol Biol. 2006;357: 373–386. doi:10.1016/j.jmb.2006.01.007

57. Dror RO, Jensen MØ, Borhani DW, Shaw DE. Exploring atomic resolution physiology on a femtosecond to millisecond timescale using molecular dynamics simulations. J Gen Physiol. 2010;135: 555–562. doi:10.1085/jgp.200910373

58. Jorgensen WL, Chandrasekhar J, Madura JD, Impey RW, Klein ML. Comparison of simple potential functions for simulating liquid water. J Chem Phys. 1983;79: 926–935. doi:10.1063/1.445869

59. Darden T, York D, Pedersen L. Particle mesh Ewald: An *N* ·log(*N*) method for Ewald sums in large systems. J Chem Phys. 1993;98: 10089–10092. doi:10.1063/1.464397

60. Berendsen HJC, Postma JPM, van Gunsteren WF, DiNola A, Haak JR. Molecular dynamics with coupling to an external bath. J Chem Phys. 1984;81: 3684–3690. doi:10.1063/1.448118

61. Martyna GJ, Klein ML, Tuckerman M. Nosé–Hoover chains: The canonical ensemble via continuous dynamics. J Chem Phys. 1992;97: 2635–2643. doi:10.1063/1.463940

62. Martyna GJ, Tuckerman ME, Tobias DJ, Klein ML. Explicit reversible integrators for extended systems dynamics. Mol Phys. 1996;87: 1117–1157. doi:10.1080/00268979600100761

63. Jacobson MP, Pincus DL, Rapp CS, Day TJF, Honig B, Shaw DE, et al. A hierarchical approach to all-atom protein loop prediction. Proteins Struct Funct Bioinforma. 2004;55: 351–367. doi:10.1002/prot.10613

